# Detecting Special Genes in Marine Symbionts: A Phylogeny-Deviation Approach Identifies Recombination-Enhancing Variants

**DOI:** 10.1101/2024.10.31.621220

**Authors:** Xiaolin Zhou, Weijie Li, Siya Xie, Kang Xia, Mei Xiao, Xiaojing Yang, Zhiyuan Li

**Affiliations:** Tsinghua-Peking Joint Center for Life Sciences, School of Life Sciences, Tsinghua University, Beijing, China; Peking University Chengdu Academy for Advanced Interdisciplinary Biotechnologies, Chengdu, China; Khoury College of Computer Sciences, Northeastern University; Peking-Tsinghua Joint Center for Life Sciences, Academy for Advanced Interdisciplinary Studies, Peking University, Beijing, China; Center for Quantitative Biology, Academy for Advanced Interdisciplinary Studies, Peking University, 100871 Beijing, China

## Abstract

Different microbes possess special traits that enable adaptation to their specific niches, often through specialized genes. Identifying such species-special genes provides insights into microbial physiology and offers new tools for bioengineering. However, beyond the search for orphan genes, there are few methods for detecting genes that are homologous to those in other species yet contain unusual functional regions. In investigating the extreme recombination frequency of the marine symbiont bacterium *Ca. E. kahalalidifaciens*, we developed a computational approach to identify vairants that substantially deviate from phylogenetic expectations. This unbiased strategy successfully identified multiple Holliday junctions-related genes in *Ca. E. kahalalidifaciens* as candidates responsible for its exceptional recombination capacity. Heterologous expression of these gene variants in *Escherichia coli* significantly enhanced recombination efficiency. These high-efficient variants offer insights into improving tools for genetic manipulations, and our gene-identification approach can be applied broadly for microbial genome mining.

## Introduction

In the quest for survival, each microorganism exhibits its unique abilities to adapt to its niche [1]. For example, *Deinococcus radiodurans* can withstand extreme levels of radiation through efficient DNA repair mechanisms [2], while *Pseudomonas syringae* produces ice-nucleating proteins that promote frost damage on plants to facilitate colonization [3]. Another remarkable example comes from a marine symbiont, where the bacterium *Ca. E. kahalalidifaciens* lives intracellularly to marine alga *Bryopsis* sp. and provides chemical defense for its host [4]. In *Ca. E. kahalalidifaciens*, twenty biosynthetic gene clusters exchange genetic material at an exceptionally high frequency—orders of magnitude higher than in other known species [5]. Such intensive recombination constitutes a mode of “diversifying evolution”, generating chemical diversity in the host’s defensive compounds. Given the demands of such unique cellular functions, it is reasonable to expect that special genes underlie these processes. A key question remains: How shall we identify gene sets linked to each microbe’s special physiology, such as the extreme recombination frequency observed in *Ca. E. kahalalidifaciens*?

Identifying genes linked to such specialized functions is challenging. Orphan genes— without homologs in other lineages—represent one class of species-special genes [6]. However, their uniqueness often complicates functional studies, requiring laborious experimental validation [7]. Moreover, many special traits rely on variants of shared homologous, not orphan genes [8]. For example, *Sirt6* is involved in DNA damage repair and is widely distributed at the eukaryote level, while rodents with *Sirt6* variants divergent at the C-terminal exhibit substantial longevity [9]. Another example is the housekeeping *rpoB* gene encoding bacterial RNA polymerase, where certain mutations help the strain to evade the suppression of antibiotics rifampin [10]. In primates, systematic methods have been developed to investigate the Human Accelerated Regions (HAR), revealing sequences that likely contribute to human-specific traits [11]. However, in the vast genomes of microbes, there are presently relatively few comparable methods for identifying the special genes that give a bacterium its unique trait [12].

The search for such variants is complicated by microbial evolution. Frequent mutations generate extensive variability, making it hard to pinpoint gene variants linked to special traits [13, 14]. Meanwhile, phylogenetic relationships shape the overall pattern of genetic variations, with recently diverged species sharing higher sequence similarity than distant relatives [15, 16]. Thus, we can establish phylogeny-based expectations for sequence similarity within homologous groups. Variants that deviate significantly from these expectations are prime candidates for species-special genes, likely responsible for unique traits in specific organisms.

Here, we introduce a computational strategy to identify species-special (SS) genes by detecting sequences that deviate from phylogenetic expectations. Applying this method to cEK uncovered the Holliday junctions-related machinearie, including *recG, ruvA/B/C*, and *ligA* as candidates contributing to its exceptional recombination. Structural analysis and selection pressure assessments revealed key mutations in DNA-binding and interaction domains. Heterologous expression of these variants in *Escherichia coli* significantly enhanced recombination efficiency, offering potential directions for genetic engineering. This pipeline provides a generalizable framework for identifying functional variants across microbial genomes.

## Method

### Data collection and reformatting

Complete genomes from 222 species of the *Flavobacteriaceae* family were downloaded from the NCBI Genome database in GenBank format. Twenty-six genomes were excluded due to the absence of 16S rRNA annotations, leaving a total of 196 species for analysis. The genome of *Ca. E. kahalalidifaciens* were obtained from Zan’s work [4].

Gene information from each genome was parsed into multiple fields, including annotations, nucleotide sequences, and amino acid sequences. Genes with suspicious characteristics were removed from further analysis; this included those annotated as pseudogenes, located in ambiguous genomic regions, not starting with methionine, or exhibiting significant mismatch between amino acid and nucleotide translations.

16S ribosomal RNA sequences were extracted from each genome file to indicate phylogenetic relationships. In instances where multiple 16S rRNA sequences were detected within a single genome, the sequence with the least mean distance to the others was selected as the representative sequence for that species.

### Orthologous group identification and gene filtering

To identify orthologues and analyze their functions, *Escherichia coli* was used as the reference strain. The GenBank file for *E. coli* strain K-12 (accession NC_000913) was downloaded from the NCBI database, yielding a total of 4,242 protein-coding genes. The program DIAMOND v2.0.5.143 was utilized to infer homology among genes across the 197 *Flavobacteriaceae* species. When multiple sequences from the same species were assigned to the same orthologous group, the sequence closest to the reference in *E. coli* was chosen as the representative sequence. Among all orthologous groups, 619 groups contained sequences from *Ca. E. kahalalidifaciens*, which were then assigned corresponding names and IDs from *E. coli* for further analysis.

### Multiple sequence alignment and calculation of sequence distances

Multiple sequence alignments for each of the 619 orthologous groups and the 16S rRNA sequences were constructed using the built-in MUSCLE algorithm in MEGA-CC v. 10.1.8, with the Neighbor-Joining method employed for clustering throughout all iterations. Based on these alignments, pairwise distance matrices for each of the 619 orthologous groups and the 16S rRNA sequences were calculated under the Jones-Taylor-Thornton (JTT) model, with a Gamma distribution with a parameter value of 1.

The amino acid sequence distance between species *a* and *b* in the *g*-th orthologous group is denoted as *O*_*g*_(*a, b*). The 16S rRNA nucleotide sequence distance between species *a* and species *b* is denoted as *Phy*(*a, b*), representing the phylogenetic difference. Assuming there are *N* species, then both matrices are *N* × *N* in size. To avoid bias induced by differences in the evolutionary rates between ortholog groups, both distances were centralized and scaled to the unit variance.

### Scoring orthologous genes in given species by deviations from phylogeny

To identify special genes, for each orthologous group *g*, we compared its amino acid sequence distances *O*_*g*_ against the 16S distances *Phy* between all pairs of species. The protein sequence divergence is represented by *O*_*g*_, whereas the phylogenetic distance is represented by *Phy*. Using *Phy* as the x-value and *O*_*g*_ as the y-value, we established a “distance space” in which each data point as a two-element vector:

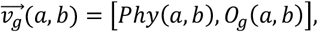

between species *a* and species *b* (total species number *N*). In this distance space, protein sequence divergence *O*_*g*_ tends to scale with phylogenetic distance *Phy* by default. We defined a special score (SS score) based on the Silhouette index [17] to quantify deviation from this default trend, as described below:

A given species *c* divides all data points 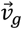 in the distance space into two clusters: the first cluster *C*_*c*_ containing points that are related to species *c* (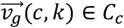, for *k* = 1 … *N*), and the second cluster *C*_other_ containing points that are unrelated to species 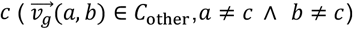. The Silhouette coefficient quantifies how well a data point in one cluster separates from points in other clusters [17]. To calculate the Silhouette coefficient, we first computed the Euclidean distance *d*_*g*_(*i, j*) between each pair *i, j* of 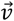 in the distance space:

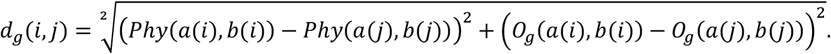

Then, for the *i*-th point in the first cluster *C*_*c*_, let *h*_*c,g*_(*i*) represent the mean Euclidean distance between this point and all other points in *C*_*c*_:

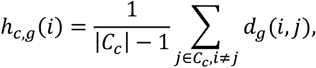

where |*C*_*c*_| is the size of number of points belonging to the first cluster.

And we let *f*(*i*) represent the mean distance of the *i*-th point in the first cluster *C*_*c*_ to all points in the second cluster *C*_other_:

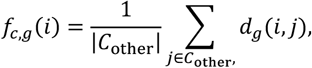

where |*C*_other_| is the size of number of points belonging to the second cluster.

The Silhouette coefficient of the *i*-th point in the first cluster *C*_1_, by definition, [17], is calculated as:

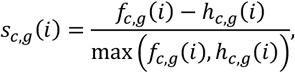

with *s*_*c,g*_(*i*) ranges from -1 to 1. A high value indicates that the *i*-th point is well matched to the first cluster and clearly distinguishable from the second cluster. Therefore, the averaged Silhouette coefficient for all points in the first cluster *C*_*c*_ reflects how much the protein-phylogeny relationships for species *c* deviate from these in all other species. We defined it as the *SSscore*(*c, g*) for species *c* in the ortholog gene *g*:

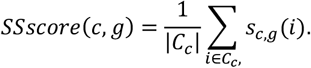

This score indicates how special this variant in the orthologous group *g* is for the species *c*, compared with the default protein-phylogeny relationship.

### Pathway enrichment analysis

As a proof of concept, we ranked the SS scores for all orthologous groups in *Ca. E. kahalalidifaciens* from highest to lowest. The top 100 ranked genes, identified by their *E. coli* names, were subjected to pathway enrichment analysis using the STRING online server.

### Motifs and structural analysis

For the gene of interest, we employed MEME Suite to detect sequence motifs within its orthologous group. PDB files for the 3D structure of *E. coli* RuvA tetramer were downloaded from the NCBI PDB database, and the 3D structure of RuvA in *Ca. E. kahalalidifaciens* was predicted using the RaptorX online server.

### Strains and culture medium

All cloning and testing experiments were conducted using *Escherichia coli* K-12 sub strain MG1655. Cells were grown in LB medium (10 g/L tryptone, 5 g/L yeast extract, and 10 g/L NaCl). For agar plates, 15 g/L agar was added to the medium. To select and maintain plasmids, antibiotics were used at concentrations of either 100 µg/mL ampicillin or 50 µg/mL kanamycin. All chemicals used in this study were purchased from Sigma–Aldrich unless otherwise stated.

### Gene Codon and Expression Optimization

To optimize the simultaneous overexpression of recombination-related proteins, including RecA, LigA, RuvA, RuvB, RuvC, and RecG, in *E. coli* MG1655, we first performed codon optimization tailored to this specific expression host. The ribosome binding sites (RBS) for each protein were subsequently calculated based on the optimized codon sequences (Table 1) using the De Novo DNA platform (https://www.denovodna.com/). The genes were then arranged in order of their predicted endogenous expression levels in *E. coli*, from low to high, according to data from EcoCyc (http://ecocyc.org). These sequences were synthesized by GeneWiz and BGI for further experimental analysis.

**Table 1.**
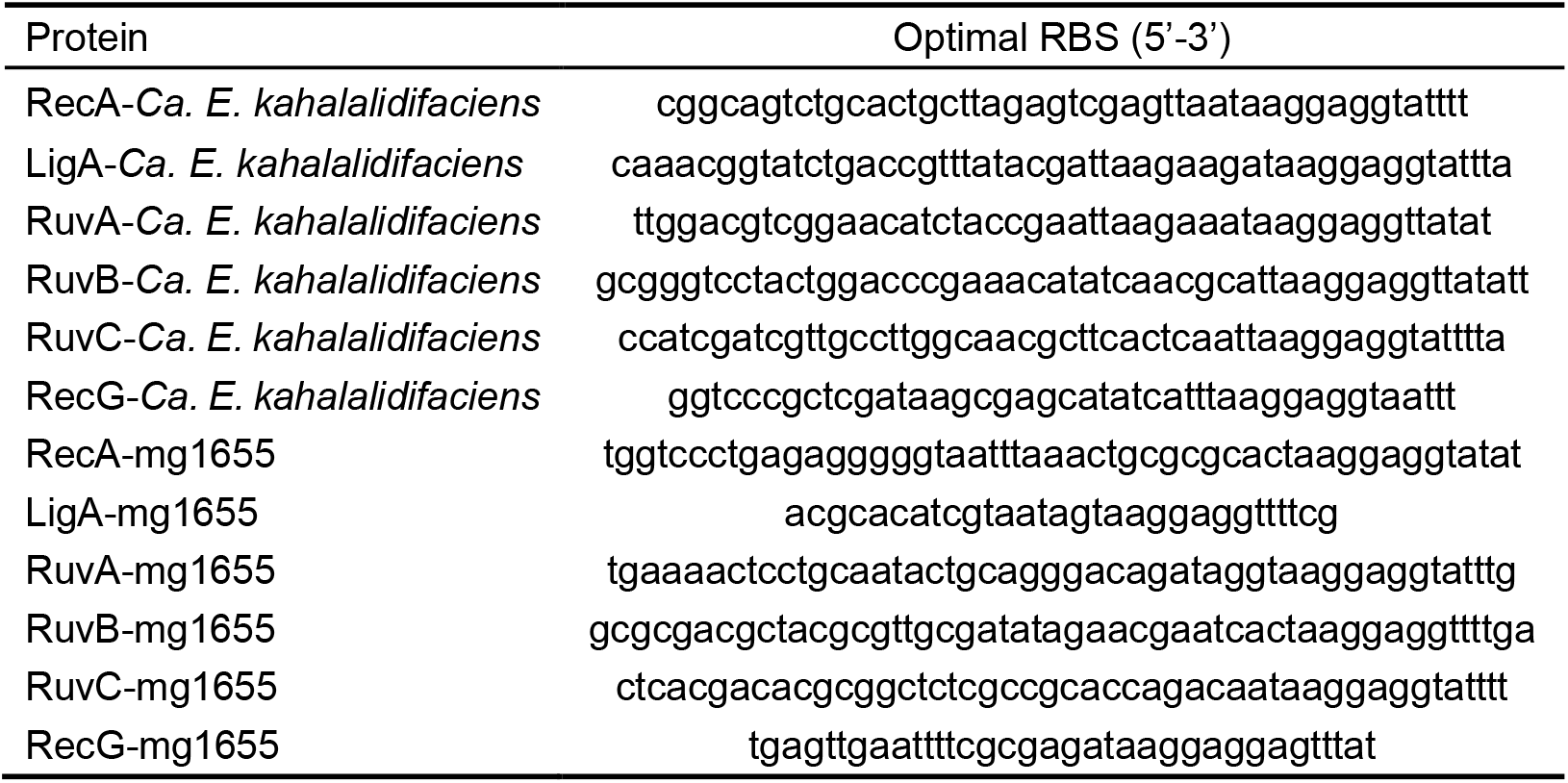
The calculated optimal RBS used in this study.

### Plasmid construction

To enable controlled overexpression of the target proteins, we utilized the L-arabinose-inducible araBAD promoter. Given the potential metabolic burden associated with overexpression, we opted for a medium-copy vector, using the p15A origin (∼15-30 copies per cell). A Kanamycin resistance marker was selected for the expression vector to ensure compatibility with the Ampicillin resistance in the homologous recombination assay kit. A complete list of primers used is provided in Table 2.

**Table 2.**
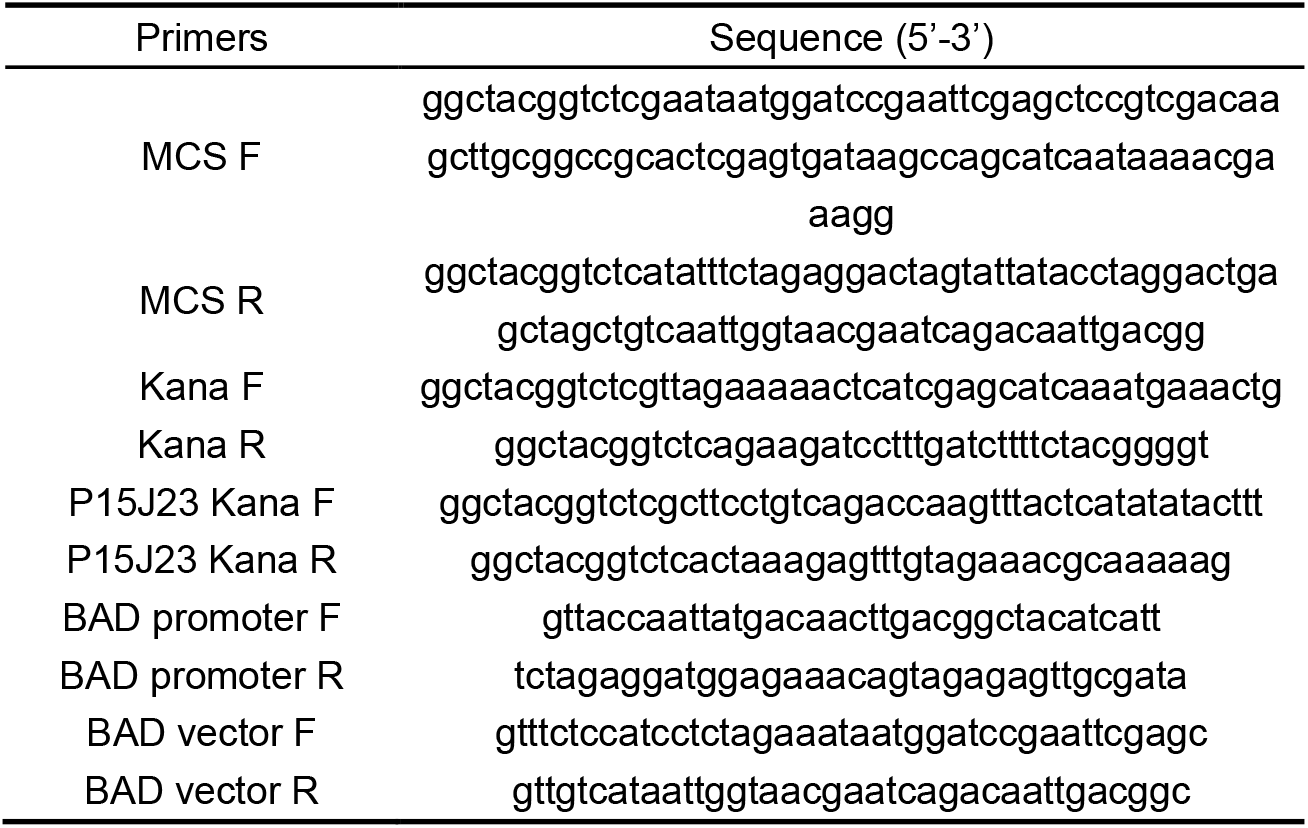
The primers used in this experiment.

Initially, we amplified a linear backbone (P15J23A) from the P4218 plasmid, kindly provided by ZhiSun, using the MCS F and MCS R primers. The P15J23A plasmid backbone included the p15A origin, Ampicillin resistance, the constitutive promoter J23119, a multi-cloning site (MCS), and a strong terminator. The linearized backbone was circularized using the Golden Gate reaction.

Next, we amplified the Kanamycin resistance gene from the pET-28a plasmid using Kana F and Kana R primers. After DpnI digestion and purification, we seamlessly cloned the backbone and the Kanamycin fragment via Golden Gate assembly, generating the p15J23K vector.

Subsequently, the araBAD promoter was amplified from the pCas plasmid using BAD promoter F and BAD promoter R primers. This fragment was assembled with a linearized vector fragment, derived from p15J23K using BAD vector F and BAD vector R primers, via Gibson assembly, yielding the P15BAD vector.

Finally, the synthesized overexpression fragments for recombinase proteins from both the *Ca. E. kahalalidifaciens* and *E. coli* MG1655 strains were inserted into the P15BAD vector through double-enzyme digestion using BamHI and HindIII, resulting in the vectors p15BAD-*Ca. E. kahalalidifaciens* and p15BAD-mg1655. The successful construction of the plasmids was confirmed through sequencing.

### Recombination system introduction

For the homologous recombination assay, we utilized the Homologous Recombination Assay Kit (Norgen Biotek, Product #35600, Canada). The plasmids p15BAD-*Ca. E. kahalalidifaciens* and p15BAD-mg1655 were first electroporated into *E. coli* MG1655 cells. Fresh competent cells carrying the overexpression plasmids were prepared the following day through repeated procedures. L-arabinose was added to a final concentration of 10 mM one hour before cell harvesting to induce the expression of recombination-related proteins.

Afterward, 25 ng of each DL-1 and DL-2 plasmids (from the assay kit) were co-transformed into the prepared competent cells. The transformation was performed at a 1:100 dilution into a liquid medium containing both Kanamycin and Ampicillin, followed by overnight culture. All transformations were done in triplicate, with MG1655 cells without the overexpression plasmid serving as the negative control.

### Quantitative analysis of recombination

Plasmid DNA was extracted from the cultures and adjusted to 20 ng/μL for further qPCR analysis. This analysis aimed to quantify the recombination rate by amplifying plasmids with both universal and assay-specific primers. Universal primers targeted all plasmids (DL-1, DL-2, and the recombined positive plasmid), while assay-specific primers amplified only the recombined plasmid. Recombination rates were calculated based on CT values obtained during qPCR with the following formula:

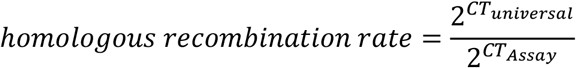

The relative recombination efficiency can then be determined based on the number of amplification cycles required to reach the threshold value for both universal and specific targets.

## Result

### Data Overview and SS Score Calculation

We analyzed 197 complete genomes from the *Flavobacteriaceae* family, including *Ca. E. kahalalidifaciens* (cEK) (Fig. 2A). Across these genomes, we identified 619 orthologous gene groups present in cEK (Fig. 2B). The genomes exhibit similar sizes and gene counts, as shown in Figures 2C and 2D, ensuring a consistent dataset for comparative analysis.

**Figure 1.**
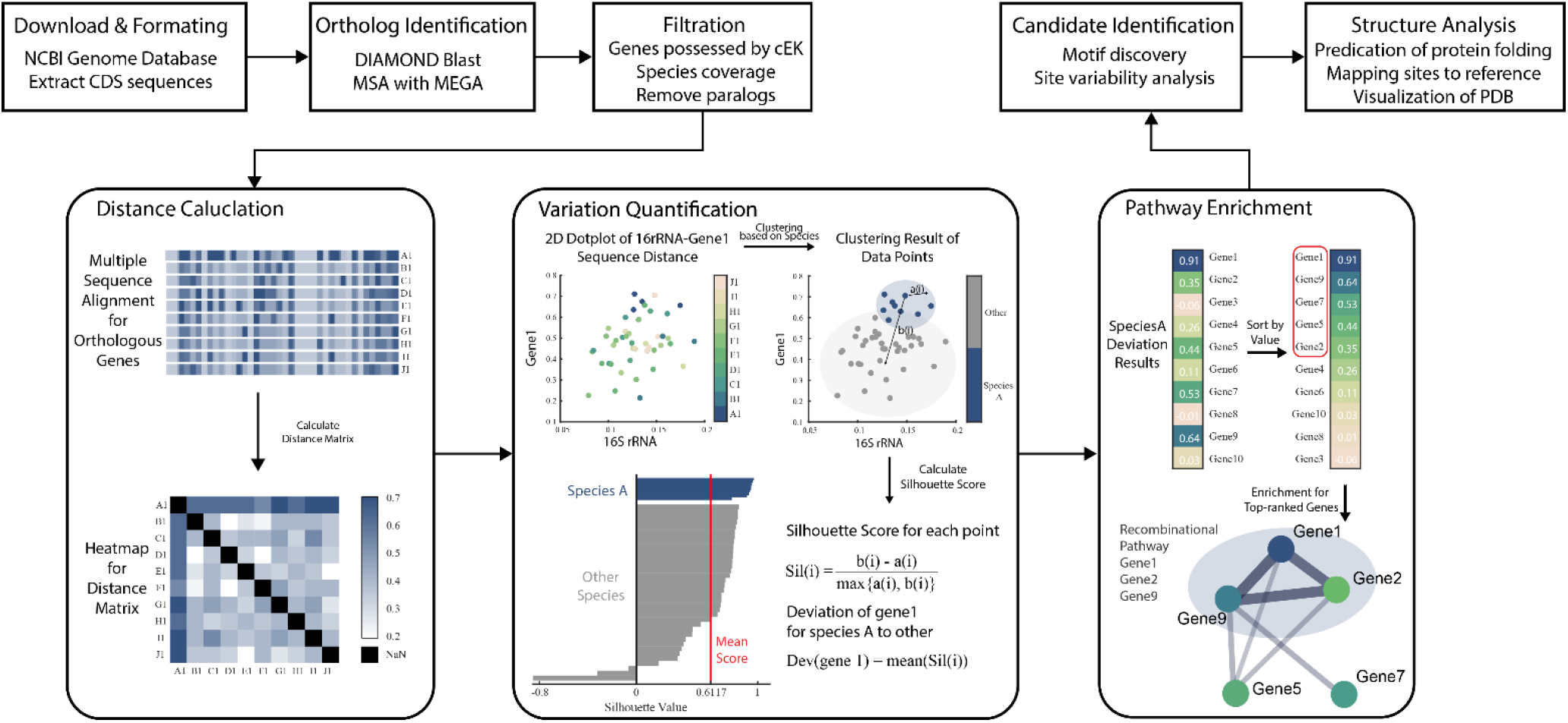
Method overview: the workflow in identifying special genes for a species.

**Figure 2.**
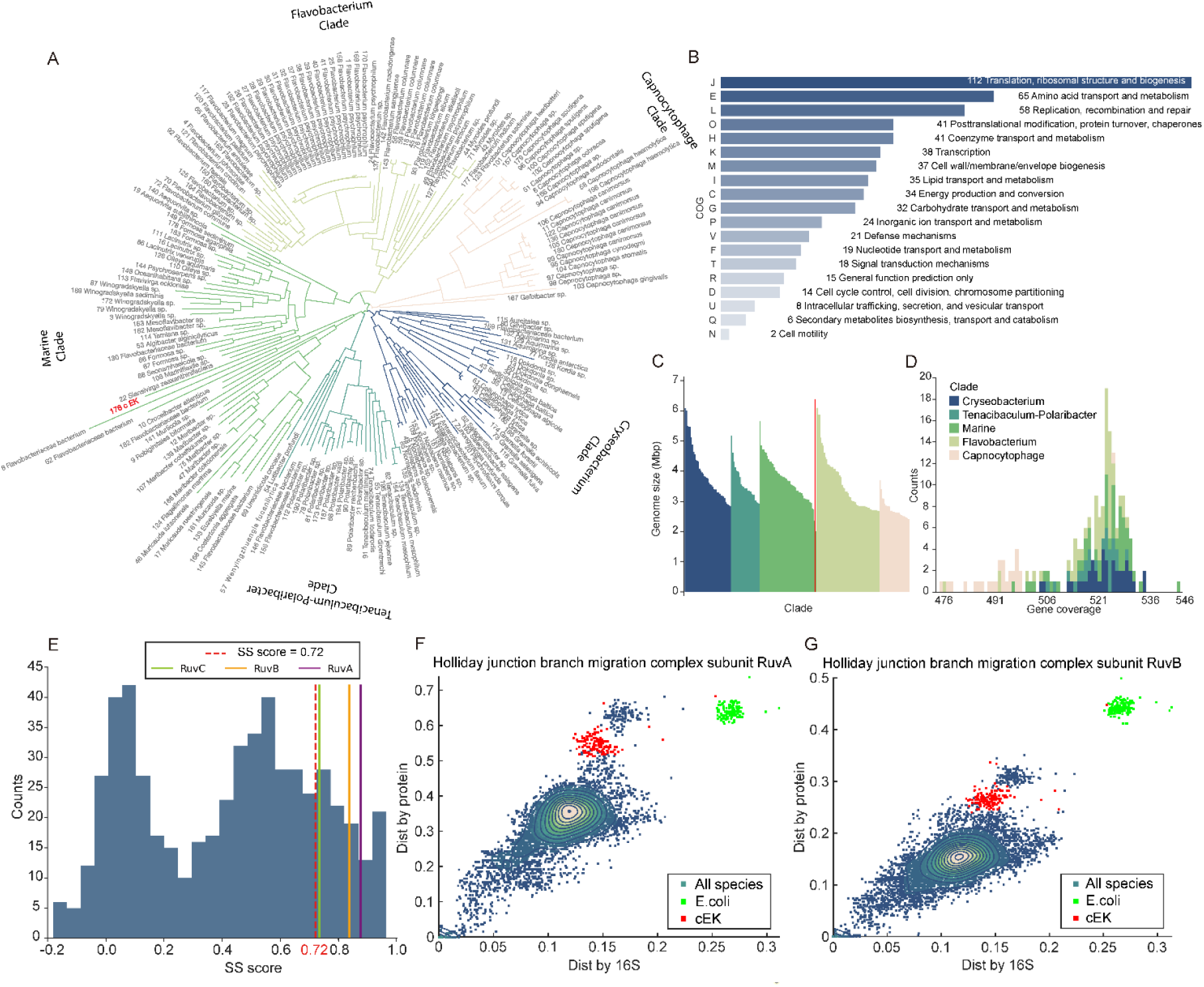
Overview of genomic data and SS score calculation. (A) Phylogenetic tree based on 16S rRNA sequences from 197 species, with *Ca. E. kahalalidifaciens* (cEK) highlighted in red. (B) Functional distribution of the 619 orthologous gene groups, categorized by COG classification. (C) Genome size distribution of the species shown in (A), with colors corresponding to their clades. (D) Gene coverage across the 619 orthologous groups within each genome, colored by clade as in (A). (E) SS score distribution for the orthologous gene groups in *Ca. E. kahalalidifaciens*. The red dashed line indicates the threshold (z-score = 1). (F, G) Comparison between pairwise protein sequence distances and 16S rRNA distances for the Holliday junction branch migration complex subunits RuvA (F) and RuvB (G). Points in red represent cEK, green points represent *E. coli*, and blue points represent all other species.

To quantify how much a gene’s protein-coding sequence deviates from expected phylogenetic patterns, we developed the “SS score” (species-special score). This metric, ranging from -1 to 1, measures the extent of deviation, with higher values indicating greater divergence from the typical phylogenetic relationship (see Methods for details).

We calculated SS scores for all 619 orthologous gene groups in cEK, yielding a distribution centered around 0.5 (Fig. 2E). To illustrate the concept and application of SS scores, we examined two key components of the Holliday junction branch migration complex: RuvA and RuvB. Figures 2F and 2G highlight the relationship between 16S rRNA-based phylogenetic distances (x-axis) and the corresponding distances in protein-coding sequences (y-axis). For most species, the distances align closely along the diagonal, reflecting the expected evolutionary relationship. However, the red points representing cEK form distinct clusters, deviating significantly from the general trend. These outliers suggest that RuvA and RuvB have undergone sequence modifications specific to cEK, potentially contributing to its unique recombination abilities.

### Unbiased search revealed that both metabolic pathways and homologous recombination processes are highly specialized in *Ca. E. kahalalidifaciens*

We selected the top 100 genes for enrichment analysis, focusing on Cellular Components and Biological Processes. The Cellular Component analysis (Table 3) identified the Holliday junction resolvase complex as highly enriched with significant confidence. In parallel, the Biological Process analysis (Table 4) showed a strong enrichment of nucleotide metabolism and translation regulation pathways, reflecting the role of cEK as an intracellular symbiont. Additionally, the recombinational repair process emerged with a high enrichment strength and a low false discovery rate (FDR), suggesting it plays a crucial role in the bacterium’s physiology.

**Table 3.** Cellular component enrichment with the false discovery rate lower than 0.01 and the strength greater than 0.75.

| Cellular Component |  |  |  |  |
| --- | --- | --- | --- | --- |
| GO-term | Description | Count in network | Strength | False discovery rate |
| GO: 0048476 | Holiday junction resolvase complex | 3 of 6 | 1.35 | 0.0072 |
| GO: 0022625 | Cytosolic large ribosomal subunit | 6 of 32 | 0.92 | 0.0021 |

**Table 4:**
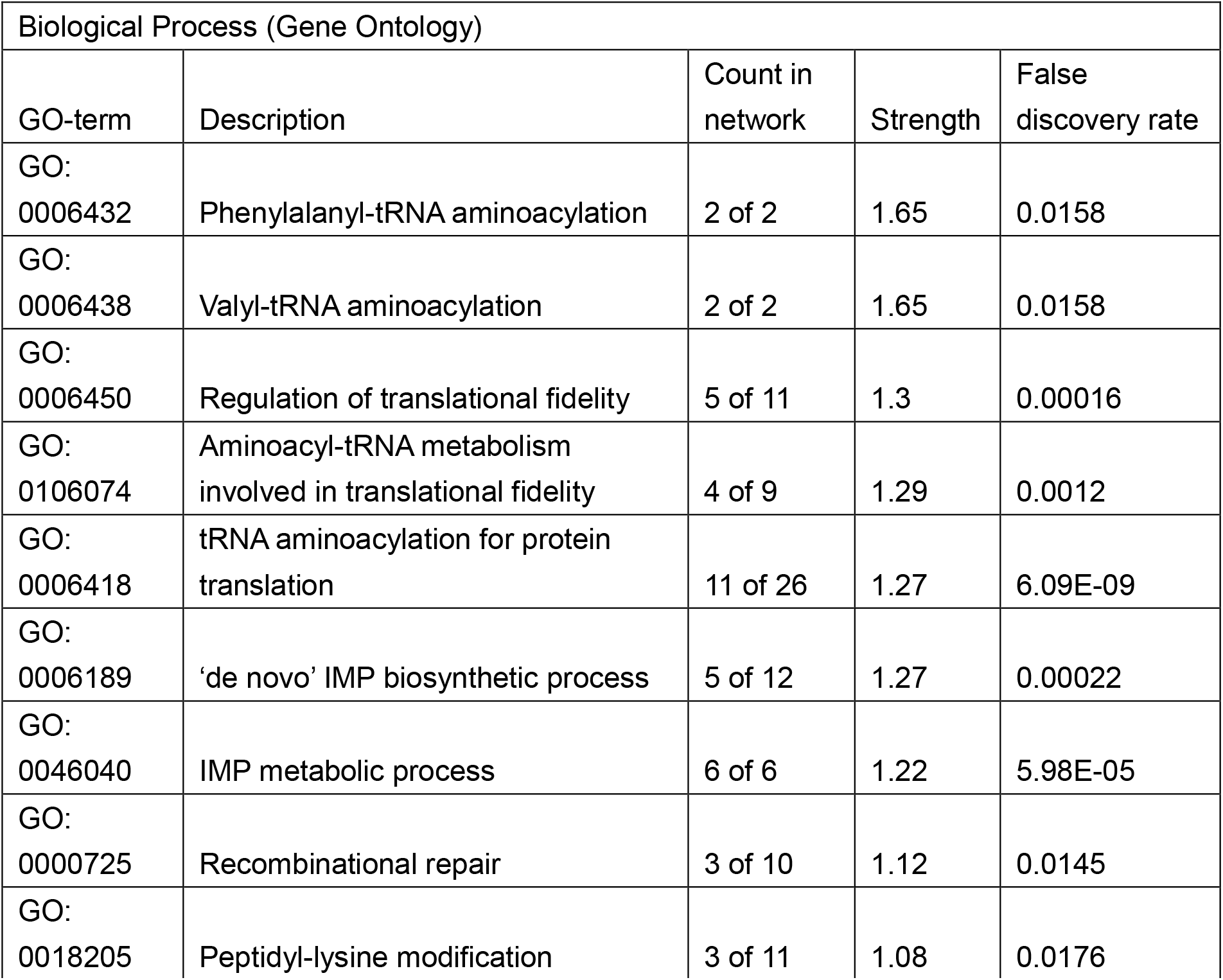
Biological Process enrichment with the false discovery rate lower than 0.02 and the strength greater than 1.

homologous recombination is an energy-consuming and high-fidelity process that demands cellular resources [18, 19]. In most microorganisms, the homologous recombination pathways are conserved [19, 20]. For the model organism *E. coli*, two general pathways of homologous recombination have been identified. One pathway operates at double-stranded DNA molecules using RecBCD, followed by RecA and either the RuvABC complex or RecG [21]. Another pathway utilizes RecF, RecO, and RecR to deal with the single-strand break, followed by RecA, then either RuvABC or RecG. In both pathways, RecA, RuvABC complex, RecG, and at last DNA ligase (to ligate DNA) are needed [22].

We further analyzed recombination-related genes with SS scores above 0.72 (z-score ≥ 1) to identify candidates special to cEK (Fig. 3A). The analysis pinpointed several key members of the Holliday junction resolvase complex: RuvA, RuvB, and RuvC, with SS scores of 0.88, 0.84, and 0.73, respectively. LigA, a NAD-dependent DNA ligase, also showed a high SS score of 0.77. The distribution of enrichment strengths (Fig. 3B) confirms the high significance of recombinational repair, with the Holliday junction resolvase complex marked by strong enrichment in the Cellular Component category. In contrast, members of the Rec family displayed lower scores, with RecG at 0.71, RecA at 0.43, and RecQ at 0.01, indicating they may play less specialized roles in this species. Interestingly, the replication-associated recombination protein RarA, with an SS score of 0.80, also emerged as a candidate, although its precise role in DNA repair remains under investigation [23]. These results suggest that the recombination machinery of cEK may have undergone specific adaptations, possibly contributing to its enhanced recombination capacity.

**Figure 3.**
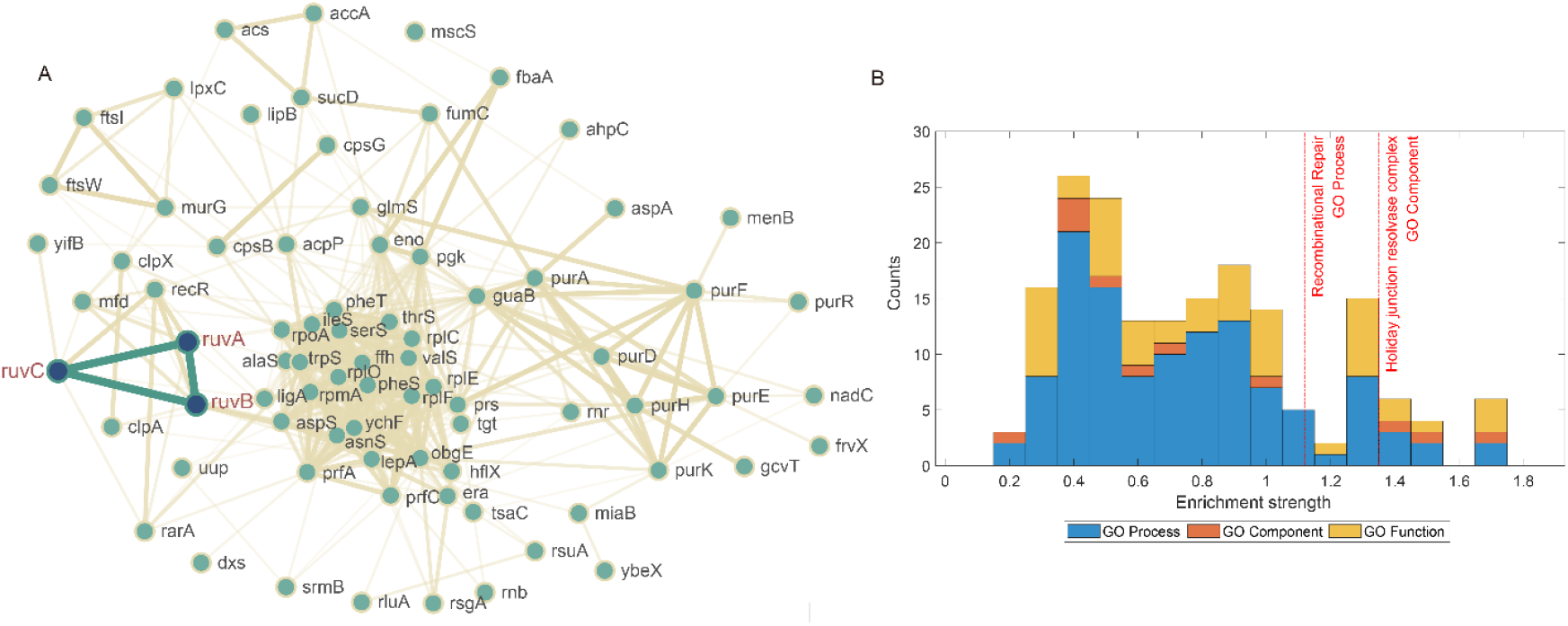
Gene ontology enrichment analysis for *Ca. E. kahalalidifaciens*-special genes. (A) Gene interaction network generated by STRING, with *ruvA, ruvB*, and *ruvC* highlighted in red. (B) Statistics of gene enrichment. The enrichment strength is log10(observed / expected).

### Sequence and Structural Analysis of *Ca. E. kahalalidifaciens*-specific Recombination-related Genes

Given that our unbiased SS score analysis revealed several recombination-related genes to be highly specialized in *Ca. E. kahalalidifaciens* (cEK), we conducted a detailed investigation of six key genes: *ruvA, ruvB, ruvC, recA, recG*, and *ligA*.

To assess evolutionary pressures on these proteins, we calculated their nonsynonymous/synonymous (dN/dS) mutation ratios, comparing cEK to 196 other *Flavobacteriaceae* species. As shown in Figure 4G, *ruvA* exhibited the highest average dN/dS ratio (1.35), followed by *recG* (1.22), while *recA* showed the lowest ratio. We further examined the dN/dS values and deviation scores across each protein’s sequence to identify regions subject to positive selection (Fig. 4A-F).

**Figure 4.**
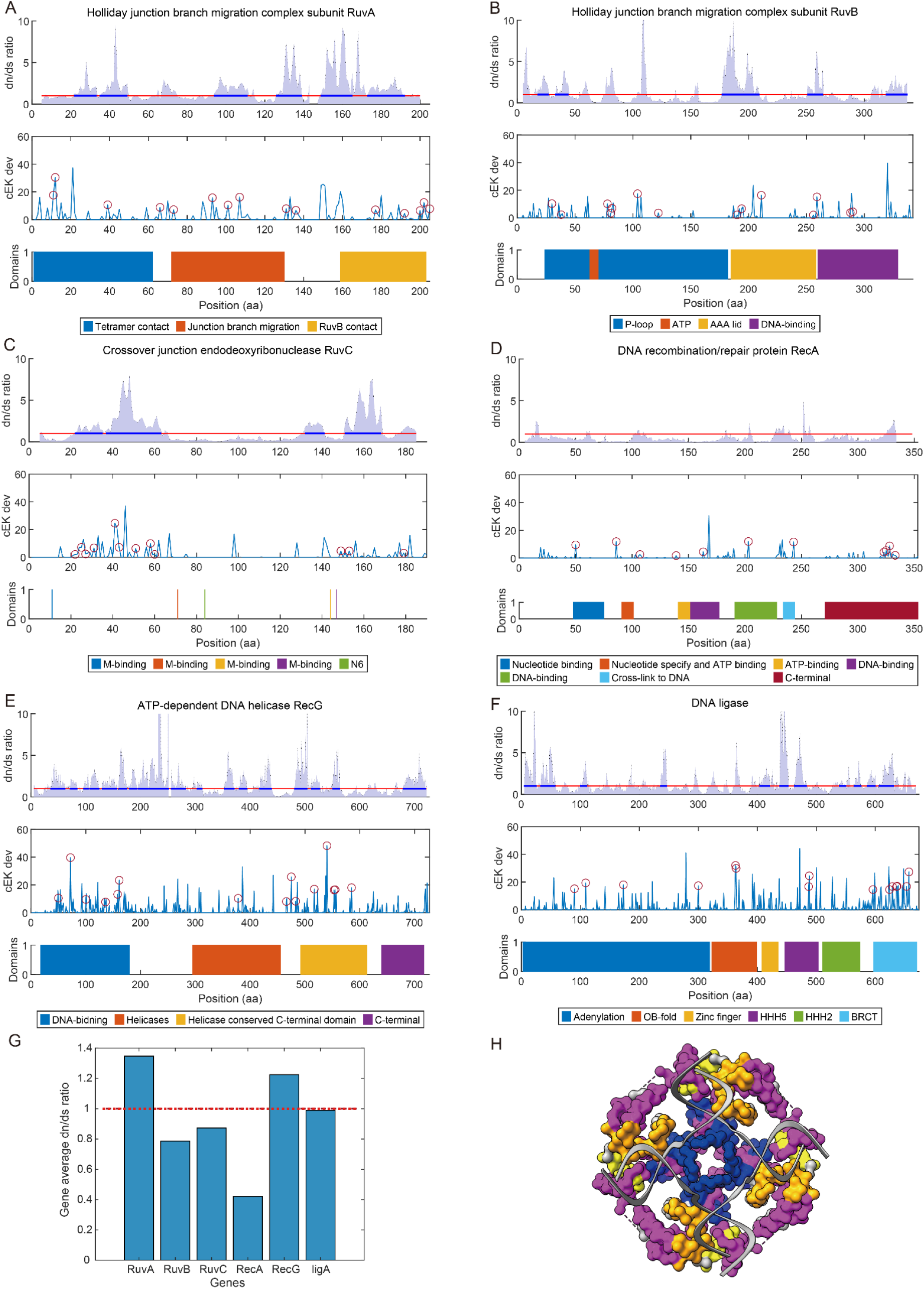
Sequence and structural analysis of *Ca. E. kahalalidifaciens*-specific recombination-related proteins. (A-F) Analysis of six recombination-related proteins: RuvA, RuvB, RuvC, RecA, RecG, and LigA. The top panel in each subplot displays the non-synonymous/synonymous mutation ratio (dN/dS) per codon site for *Ca. E. kahalalidifaciens* (cEK) compared to 196 other *Flavobacteriaceae* species. The red line indicates the threshold of dN/dS = 1, with regions above this threshold for at least 10 consecutive sites highlighted in blue. The middle panel shows the amino acid distance per site between cEK and other species, with red circles marking the 15 sites with the highest deviation that also differ from the *E. coli* sequence. The bottom panel provides functional site annotations and domain structures for each protein. (G) Average dN/dS ratios for the six proteins in cEK compared to other *Flavobacteriaceae* species. The red dashed line marks the neutral evolution threshold (dN/dS = 1). (H) Structural model of RuvA bound to a Holliday junction. The RuvA tetramer is colored according to the functional domains: blue indicates tetramer binding regions, orange denotes junction branch migration domains, yellow highlights RuvB contact sites, and the remaining regions are shown in grey. Regions with continuously elevated dN/dS values (>1) are highlighted in pink, indicating potential adaptive changes.

*recA*, which initiates recombination by facilitating single-strand DNA invasion [24], displayed low SS scores (0.43) and did not show signs of significant positive selection in cEK (Fig. 4D). This suggests that *recA* has not diverged considerably from its phylogenetic relatives in cEK.

The Holliday junction, formed after RecA-mediated strand invasion, is processed by a complex consisting of a RuvA tetramer, two RuvB hexameric rings, and a RuvC dimer. Together, these proteins facilitate junction migration and resolution [25]. Within this complex, RuvA binds to DNA and senses structure, RuvB provides ATP-driven energy for migration, and RuvC acts as an endonuclease to resolve the junction [26].

Our analysis revealed that domains involved in protein-protein interactions within the RuvA/B/C complex are under strong positive selection in cEK (Fig 4A-C). For RuvA in *Ca. E. kahalalidifaciens*, selective pressure was concentrated in three domains responsible for tetramer contact, junction migration, and RuvB interaction (Fig 4A and Fig 4H). In other species, these three regions have been reported to be highly conserved [27]. For RuvB, its N and M domains are responsible for the RuvA-RuvB interaction and RuvB hexamer assembly, while the C domain is responsible for DNA binding [28]. In our result, high positive selection pressure was observed in the RuvA-RuvB and RuvB-RuvB interaction regions (Fig 4B).The RuvC protein can both bind to DNA and, with the existence of Mg2+, catalyze sequence-specific cleavage [27]. Multiple active sites have been identified for catalysis sequence specificity, yet few studies have divided the function of RuvC into different domains as RuvA/B does. In our observation, two regions in RuvC (residues 15-28 and residues 30-55) were detected with strong positive selection in *Ca. E. kahalalidifaciens* (Fig 4C), indicating potential functional divergence.

Other than the RuvA/B/C complex, RecG, a DNA-dependent ATPase, is also capable of contacting with Holliday junction directly to drive branch migration [29]. RecG comprises multiple functional domains: the N-terminal wedge domain interacting with the DNA junction, the helicases and the helicase conserved C-terminal domain, and the C-terminal domain that forms a hook with the first domain to wrap around the extended alpha-helix [30].In our observation, all these functional domains experience high positive selection in *Ca. E. kahalalidifaciens* (Fig4 e). Of note, most *Ca. E. kahalalidifaciens*-specific mutations concentrate in the helix-forming domains known to be conserved in most microorganisms [31].

LigA, the NAD-dependent DNA ligase that catalyzes the formation of phosphodiester linkages between 5’-phosphoryl and 3’-hydroxyl groups in double-stranded DNA, functions at the last step of recombination in ligating the DNA strand [32, 33]. More importantly, it has been suggested to function in other non-homologous DNA repair pathways, including the non-homologous end-joining (NHEJ) [34] and the alternative end-joining (A-EJ) pathway [35], and offers potentials for efficient CRISPR-Cas9-assisted genetic engineering in microbes [36]. In *Ca. E. kahalalidifaciens*, the functional domains of LigA consistently exhibit a high dN/dS ratio (Fig 4F), particularly at the BRCA1 C-terminal, the integral signaling module in the DNA damage response [37]. As LigA is involved in multiple DNA damage response pathways, its *Ca. E. kahalalidifaciens*-specific mutations may bias it towards homologous recombination repair.

### *Ca. E. kahalalidifaciens* proteins significantly enhance homologous recombination in *E. coli*

Our bioinformatic analysis identified several *Ca. E. kahalalidifaciens* (cEK) proteins with high potential to enhance homologous recombination. Based on these findings, we hypothesized that the cEK recombination system has evolved unique adaptations to increase recombination efficiency, and we proceeded to validate this experimentally.

To test whether the cEK recombination proteins enhance homologous recombination in *E. coli*, we introduced the cEK recombination system into wild-type *E. coli* MG1655 and measured recombination rates. For comparison, we also introduced an additional copy of the native *E. coli* recombination system as a control. To optimize expression in *E. coli*, we adjusted codon usage and ribosome binding sites (RBS) for each cEK gene. All six genes were organized in a single operon under the control of an inducible promoter, arranged according to their endogenous expression levels in *E. coli* to facilitate efficient protein interaction (Fig. 5A). By using gradient arabinose induction, we ensured that any observed changes in recombination rates were due to the introduced cEK proteins.

**Figure 5.**
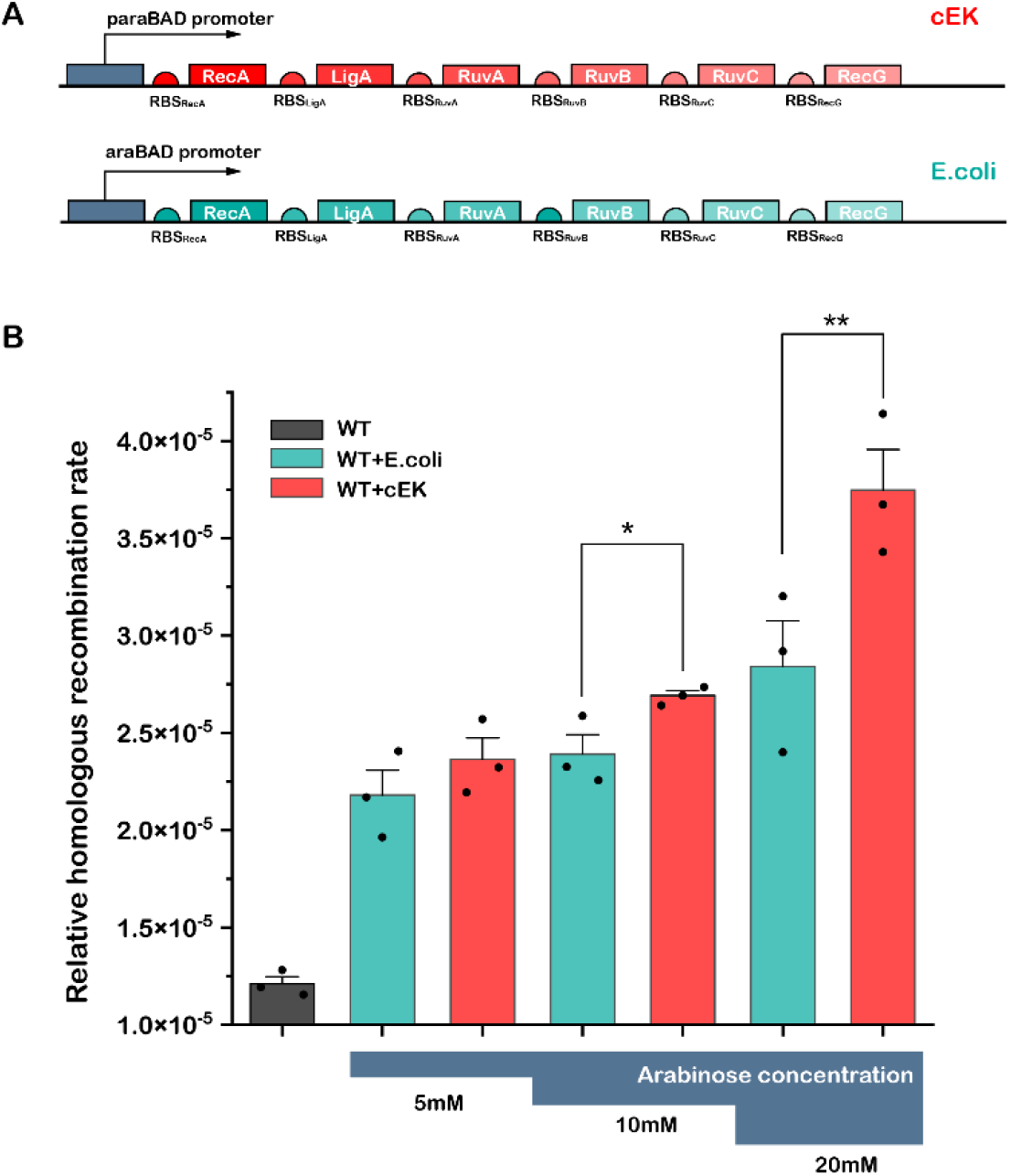
*Ca. E. kahalalidifaciens* proteins significantly enhance homologous recombination in *E. coli*. (A) Schematic representation of plasmid constructs used to express homologous recombination systems. The upper panel illustrates the homologous recombination system derived from *Ca. E. kahalalidifaciens* (cEK), while the lower panel shows the native *E. coli* system, both under the control of the arabinose-inducible paraBAD promoter. (B) Comparison of homologous recombination rates across three experimental groups: wild-type *E. coli* (gray), wild-type *E. coli* expressing the native *E. coli* recombination system (teal), and wild-type *E. coli* expressing the cEK recombination system (red). Recombinational efficiency was measured at three arabinose concentrations (5 mM, 10 mM, and 20 mM). Statistical significance is indicated as *P < 0.05 and **P < 0.01

The results showed that expression of the cEK recombination system significantly increased homologous recombination rates in *E. coli* compared to the native *E. coli* system, with the effect becoming more pronounced at higher induction levels (Fig. 5B).

## Discussion

In this study, we identified and experimentally validated the role of *Ca. E. kahalalidifaciens* (cEK) proteins in enhancing homologous recombination. Using an SS score-based bioinformatic approach, we pinpointed recombination-related genes with high potential for species-specific adaptation. Experimental validation in *E. coli* showed that the cEK recombination system significantly improved recombination rates, supporting the hypothesis that cEK’s recombination machinery has evolved for enhanced efficiency. These findings reveal unique aspects of the cEK homologous recombination pathway and suggest possible applications in genetic engineering.

Our analysis highlighted substantial positive selection in key recombination proteins RuvA, RuvB, RuvC, RecG, and LigA. Recombination has been a key factor in genetic engineering [38]. Despite that most engineering efforts in optimizing recombination efficiency have been put on RecA [39], as well as its eukaryote ortholog Rad51 [40], the conserved nature of RecA in *Flavobacteriaceae* suggests that it may maintain an essential role in initiating homologous recombination, which may be more challenging to optimize further without compromising its essential functions. In contrast, the RuvA/B/C complex in cEK appears to have undergone substantial adaptive modifications, likely enabling fine-tuning downstream processes like branch migration and junction resolution. Our finding implies that engineering efforts in recombination could benefit from focusing on the Ruv complex, as they show greater evolutionary flexibility. The distinct adaptations observed in cEK’s Ruv system could serve as a foundation for bioengineering strategies aimed at enhancing recombination efficiency in other systems.

Beyond our specific findings, this work underscores the value of the SS score as a powerful, expandable, and intuitive method for identifying species-specialized genes. It does not constrain the specific function the gene performs, just detecting anything that deviates from the phylogenetic expectation, allowing researchers to isolate candidates for specialized traits in diverse microbial genomes. The simplicity of the method also makes it accessible for broad applications across various fields, from evolutionary biology to synthetic biology, enabling the discovery of lineage-specific adaptations without prior knowledge of gene function [41].

While our study provides important insights, several limitations remain. The SS score relies on full sequence comparisons, which may miss genes with functionally significant but subtle sequence changes. Additionally, our analysis focuses on coding sequences; non-coding regulatory elements could also play a role in the evolution of specialized traits and warrant further investigation [42]. Future improvements could incorporate structural analysis and experimental validation across a broader range of organisms to refine the SS score’s predictive power. By expanding these capabilities, the SS score could become a key tool for uncovering specialized genetic adaptations across diverse lineages.

## Ethics Statement

This article does not contain any studies with human or animal subjects performed by any of the authors.

## Acknowledgments

This work was supported by the National Natural Science Foundation of China (No. 32071255, No. T2321001), and the National Key Research and Development Program of China (2021YFF1200500). LZ was supported in part by the Peking-Tsinghua Center for Life Sciences.

The experiment strain MG1655 is a kindly gift from Yiping Wang from Peking University

## Reference

1. Bleuven, C. and C.R. Landry, Molecular and cellular bases of adaptation to a changing environment in microorganisms. Proceedings of the Royal Society B, 2016. 283(1841): p. 20161458.

2. Cox, M.M. and J.R. Battista, Deinococcus radiodurans — the consummate survivor. Nature Reviews Microbiology, 2005. 3(11): p. 882–892.

3. Forbes, J., et al., Water-organizing motif continuity is critical for potent ice nucleation protein activity. Nature Communications, 2022. 13(1): p. 5019.

4. Zan, J., et al., A microbial factory for defensive kahalalides in a tripartite marine symbiosis. Science, 2019. 364(6445).

5. Li, Z., et al., Natural diversifying evolution of nonribosomal peptide synthetases in a defensive symbiont reveals nonmodular functional constraints. PNAS Nexus, 2024. 3(9): p. pgae384.

6. Johnson, B.R., Taxonomically restricted genes are fundamental to biology and evolution. Frontiers in genetics, 2018. 9: p. 407.

7. Khalturin, K., et al., More than just orphans: are taxonomically-restricted genes important in evolution? Trends in Genetics, 2009. 25(9): p. 404–413.

8. Liang, Y.T., et al., Recent advances in the characterization of essential genes and development of a database of essential genes. Imeta, 2024. 3(1): p. e157.

9. Tian, X., et al., SIRT6 Is Responsible for More Efficient DNA Double-Strand Break Repair in Long-Lived Species. Cell, 2019. 177(3): p. 622-638.e22.

10. Cutugno, L., et al., rpoB mutations conferring rifampicin-resistance affect growth, stress response and motility in Vibrio vulnificus. Microbiology, 2020. 166(12): p. 1160–1170.

11. Hubisz, M.J. and K.S. Pollard, Exploring the genesis and functions of Human Accelerated Regions sheds light on their role in human evolution. Current opinion in genetics & development, 2014. 29: p. 15–21.

12. Zoccarato, L., et al., A comparative whole-genome approach identifies bacterial traits for marine microbial interactions. Communications Biology, 2022. 5(1): p. 276.

13. Shu, W.-S. and L.-N. Huang, Microbial diversity in extreme environments. Nature Reviews Microbiology, 2022. 20(4): p. 219–235.

14. Wheeler, N.E., et al., A profile-based method for identifying functional divergence of orthologous genes in bacterial genomes. Bioinformatics, 2016. 32(23): p. 3566–3574.

15. Martiny, J.B., et al., Microbiomes in light of traits: a phylogenetic perspective. Science, 2015. 350(6261): p. aac9323.

16. Koonin, E. and M.Y. Galperin, Sequence—evolution—function: computational approaches in comparative genomics. 2002.

17. Rousseeuw, P.J., Silhouettes: a graphical aid to the interpretation and validation of cluster analysis. Journal of computational and applied mathematics, 1987. 20: p. 53–65.

18. Sagi, D., T. Tlusty, and J. Stavans, High fidelity of RecA-catalyzed recombination: a watchdog of genetic diversity. Nucleic acids research, 2006. 34(18): p. 5021–5031.

19. Rocha, E.P.C., E. Cornet, and B. Michel, Comparative and evolutionary analysis of the bacterial homologous recombination systems. PLoS genetics, 2005. 1(2): p. e15.

20. Hoff, G., et al., Implication of RuvABC and RecG in homologous recombination in Streptomyces ambofaciens. Research in microbiology, 2017. 168(1): p. 26–35.

21. Rafferty, J.B., et al., Crystal structure of DNA recombination protein RuvA and a model for its binding to the Holliday junction. Science, 1996. 274(5286): p. 415–421.

22. Cromie, G.A., J.C. Connelly, and D.R. Leach, Recombination at double-strand breaks and DNA ends: conserved mechanisms from phage to humans. Molecular cell, 2001. 8(6): p. 1163–1174.

23. Romero, H., et al., Single molecule tracking reveals functions for RarA at replication forks but also independently from replication during DNA repair in Bacillus subtilis. Scientific reports, 2019. 9(1): p. 1–13.

24. Cox, M.M., Regulation of Bacterial RecA Protein Function. Critical Reviews in Biochemistry and Molecular Biology, 2007. 42(1): p. 41–63.

25. Kowalczykowski, S.C., et al., Biochemistry of homologous recombination in Escherichia coli. Microbiological reviews, 1994. 58(3): p. 401–465.

26. West, S.C., Processing of recombination intermediates by the RuvABC proteins. Annual review of genetics, 1997. 31(1): p. 213–244.

27. Sharples, G.J., S.M. Ingleston, and R.G. Lloyd, Holliday junction processing in bacteria: insights from the evolutionary conservation of RuvABC, RecG, and RusA. Journal of bacteriology, 1999. 181(18): p. 5543–5550.

28. Ohnishi, T., et al., Structure-Function Analysis of the Three Domains of RuvB DNA Motor Protein*[boxs]. Journal of Biological Chemistry, 2005. 280(34): p. 30504–30510.

29. Whitby, M.C. and R.G. Lloyd, Targeting Holliday junctions by the RecG branch migration protein of Escherichia coli. Journal of Biological Chemistry, 1998. 273(31): p. 19729–19739.

30. Singleton, M.R., S. Scaife, and D.B. Wigley, Structural analysis of DNA replication fork reversal by RecG. Cell, 2001. 107(1): p. 79–89.

31. Rudolph, C.J., et al., Is RecG a general guardian of the bacterial genome? DNA repair, 2010. 9(3): p. 210–223.

32. Yang, Y., et al., The pathway of recombining short homologous ends in Escherichia coli revealed by the genetic study. Molecular Microbiology, 2021. 115(6): p. 1309–1322.

33. Shuman, S. and M.S. Glickman, Bacterial DNA repair by non-homologous end joining. Nature Reviews Microbiology, 2007. 5(11): p. 852–861.

34. Bhattacharyya, S., et al., Phage Mu Gam protein promotes NHEJ in concert with Escherichia coli ligase. Proceedings of the National Academy of Sciences, 2018. 115(50): p. E11614–E11622.

35. Chayot, R., et al., An end-joining repair mechanism in Escherichia coli. Proceedings of the National Academy of Sciences, 2010. 107(5): p. 2141–2146.

36. Huang, C., et al., CRISPR-Cas9-assisted native end-joining editing offers a simple strategy for efficient genetic engineering in Escherichia coli. Applied microbiology and biotechnology, 2019. 103(20): p. 8497–8509.

37. Leung, C.C.Y. and J.M. Glover, BRCT domains: easy as one, two, three. Cell cycle, 2011. 10(15): p. 2461–2470.

38. Tao, M., et al., Construction of a CRISPR-based paired-sgRNA library for chromosomal deletion of long non-coding RNAs. Quantitative Biology, 2020. 8(1): p. 31–42.

39. Kim, T., et al., Directed evolution of RecA variants with enhanced capacity for conjugational recombination. PLoS genetics, 2015. 11(6): p. e1005278.

40. Jin, Y.-Y., et al., Enhancing homology-directed repair efficiency with HDR-boosting modular ssDNA donor. Nature Communications, 2024. 15(1): p. 6843.

41. Ferrer, M., et al., Metagenomics for mining new genetic resources of microbial communities. Journal of molecular microbiology and biotechnology, 2009. 16(1-2): p. 109–123.

42. Tatman, B.P., et al., Significance of differential allelic expression in phenotypic plasticity and evolutionary potential of microbial eukaryotes. Quantitative Biology, 2021. 9(4): p. 400–410.

